# Solid state NMR characterization of wild-type and mutant GFAP intermediate filament assemblies

**DOI:** 10.64898/2026.05.15.725530

**Authors:** Kayla M. Osumi, Dylan T. Murray

## Abstract

GFAP is a type III intermediate filament primarily found within astrocytes and is known to maintain proper cell structure and mechanical strength. Mutations in GFAP are implicated in the pathology of Alexander disease, a neurodegenerative disease characterized by cytoplasmic inclusions of protein, known as Rosenthal fibers. GFAP has a typical type III intermediate filament domain structure, consisting of a highly conserved alpha-helical rod domain bracketed by an intrinsically disordered N-terminal head and C-terminal tail domains. While the general domain organization of monomeric GFAP and the assembly process for higher order quaternary structures are known, we lack an atomic resolution mechanistic understanding of GFAP assembly into mature filaments. Understanding the structure of GFAP filaments and how mutations disrupt this structure will provide vital information into how mutations produce Alexander disease pathology. As a first step towards a mechanistic description, we characterized GFAP wild type tetrameric and filamentous assemblies using solid state NMR and compared the results to those obtained from an assembly-deficient GFAP mutant. For wild-type GFAP, we observe surprisingly uniform rigid alpha helical structure and can spectroscopically resolve highly mobile intrinsically disordered regions in the filament assemblies. Wild type tetramers show increased mobility, likely arising from the head and tail domains. Mutation of the highly conserved cysteine at position 294 to serine results in an inability to form full-length filament assemblies. We show that the rigid regions of the C294S mutant assemblies largely remain structurally consistent with wild type tetrameric assemblies but differ from wild-type filament assemblies. There is an increase in highly mobile regions for the C294S mutant relative to the wild-type. Our results provide a foundation for developing solid state NMR approaches to characterize intermediate filament assembly mechanisms and the interfering effect of disease mutations.

## Introduction

Intermediate filament proteins (IFs) make up one of the three main components of the metazoan cell cytoskeleton and are one of the 100 largest gene families within humans.^1,2^ Despite this, there is a lack of high-resolution structural information regarding these biological assemblies.^3^ IFs have long been thought to mainly provide structural support and mechanical stability to cells. However, recent studies indicate that IFs also play an important role in cell organization, signaling, and motility.^1,3–5^ These IFs self-assemble, forming networks that extend from the nucleus to the plasma membrane and act as a scaffold that helps maintain cell structural integrity.^1^ Extended IF networks are also known to be involved in cell signaling and may play a role in mechanotransduction.^6,7^ IF proteins are ubiquitous and highly localized, with different IFs being predominantly expressed in certain cell types.^1^

There are six different types of IF categorized by sequence similarity. All six types of IF have similar domain organization with a highly conserved alpha-helical rod domain and intrinsically disordered N-terminal head and C-terminal tail domains (Fig. 1).^1^ Previous structural characterization using X-ray crystallography and electron paramagnetic resonance (EPR) has focused primarily upon the type III protein, vimentin.^8–11^ X-ray crystallography characterizations of fragments of the rod domain show the protein forms a parallel coiled-coil dimer, and a staggered antiparallel tetramer.^12^ The staggered antiparallel tetramer model is further corroborated through chemical cross-linking and site-directed spin labeling EPR (SDSL-EPR) experiments. SDSL-EPR experiments have examined residues within the core of the coiled-coil as well as within the intrinsically disordered head and tail domain. These experiments have shown that upon filament assembly, the head domain becomes more ordered, while the rod domain remains ridid and the tail domain retains mobility.^13–15^ The tail domain remains highly dynamic within the assemblies, however, rigid regions within the tail have also been identified.^13^

**Figure 1.**
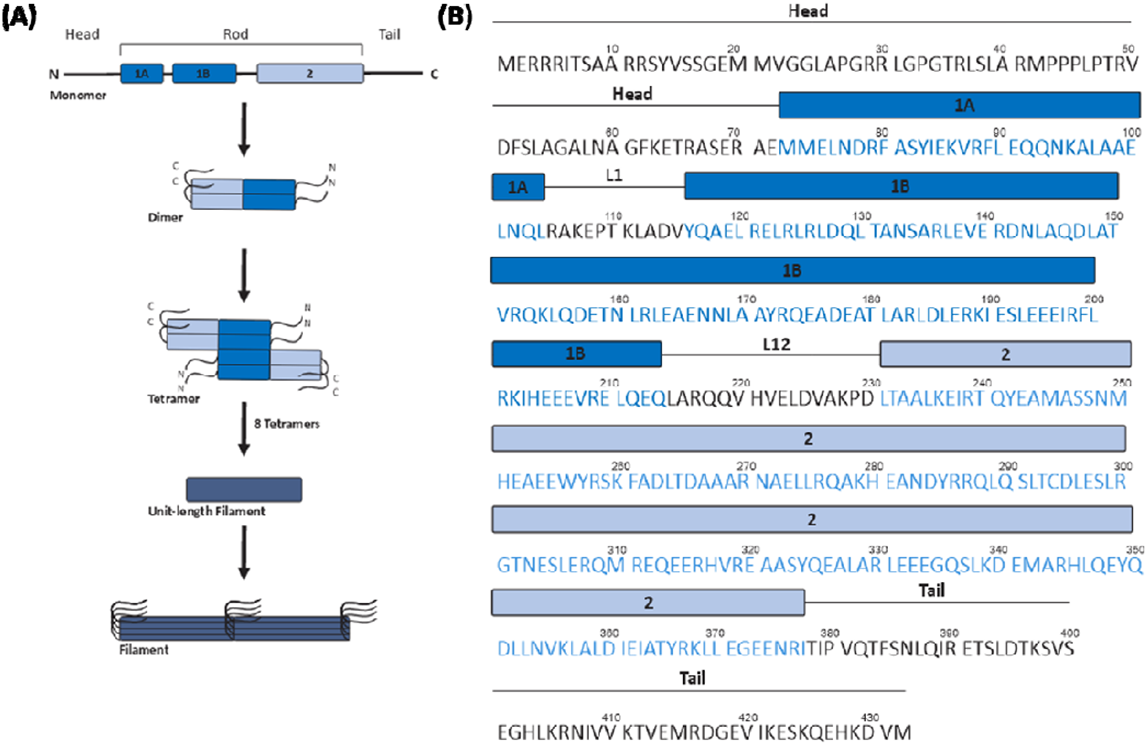
Cartoon depiction of intermediate filament assembly and structure. (A) Cartoon schematic of intermediate filament assembly. (B) GFAP amino acid sequence. Black line and text represent intrinsically disordered domains. Dark blue boxes and text represent alpha-helical coil 1 domain and light blue boxes and text represent alpha-helical coil 2 domain.

Glial Fibrillary Acidic Protein (GFAP) intermediate filament proteins are primarily found within astrocytes. GFAP filaments provide structural and mechanical strength within glial cells and enables cell motility in normal astrocyte function. After CNS injury, GFAP is upregulated as part of the process of reactive gliosis and may aid in neuroregeneration.^12^ GFAP-knockout mice have a reduced ability to handle mechanical stress and a higher mortality rate after traumatic head injury.^16^

Interestingly, it has been shown that mutation to the cysteine at position 294 in the GFAP rod domain leads to impaired and inefficient filament assembly within transfected cells.^17^ Type III intermediate filaments all contain a highly conserved cysteine located at the same position within the rod domain.^18^ It is known that a cysteine-free mutant of Vimentin forms morphologically similar filaments *in vitro* to its wild type counterpart.^18^ Within cells, the cysteine-free Vimentin filament network formation is stunted, but not nearly to the timescale and severity of cysteine-free GFAP. Cysteine-free Vimentin forms networks within hours, while it takes days for cysteine-free GFAP to do the same^17–19^. As it has been commonly assumed that intermediate filaments follow similar assembly pathways, it is surprising that this equivalent cysteine substitution impairs assembly of GFAP to such a large extent. This cysteine is thought to be important for the redox response of IF proteins, in which the cysteine is chemically modified leading to the rearrangement of the filament assemblies.^18^

Despite the importance of IFs in the cell cytoskeleton, signaling, cell motility, wound healing, and disease, there is a lack of high-resolution mechanistic understanding of how IF proteins assembly functionally and pathologically. While a moderate resolution structural model of Vimentin filaments was recently obtained^20^, a high-resolution characterization of IF proteins at various stages of assembly, disassembly, and disease will provide a mechanistic basis for IF function and IF disease treatment. In this work, we present a set of solid state nuclear magnetic resonance (NMR) measurements on wild-type and cysteine mutant GFAP mature filaments and tetrameric assemblies. Our results provide insight into the structure, stability, and flexibility of GFAP filament assemblies, and lays a foundation for the high-resolution characterization of IF protein assemblies using solid state NMR.

## Materials and Methods

### Mutant GFAP Plasmid

The C328S GFAP mutant was prepared using Gibson cloning, using two sets of forward and reverse primers containing the C328S mutation to create fragments of the plasmid. Fragment 1 was created using the primers 5’-cagagcctgacgtccgatttggagtccttg-3’ and 5’-tagctcactcattaggcaccgggatctcgacc-3’; fragment 2 was created using 5’-ggtcgagatcccggtgcctaatgagtgagcta-3’ and 5’-caaggactccaaatcggacgtcaggctctg-3’ (Integrated NDA Technologies). A polymerase chain reaction (PCR) was conducted to create the fragments, adding 1.5 ul of wild type GFAP plasmid (79.6 ng/ul—see *Protein Expression* section), 1 uL of 10 mM deoxynucleotides, 1.5 uL of dimethyl sulfoxide, 2.5 uL of 10 uM forward and reverse primer, 0.5 uL of Phusion HiFi polymerase, and 10 uL of 5x Phusion GC buffer (New England BioLabs). Autoclaved ultrapure water was added to a reaction volume of 50 uL. The reaction was carried out using a Bio-Rad T100 thermal cycler. The initial denaturation step was carried out at 98 °C for 30 seconds. Next, 30 cycles of 98 °C denaturation for 7 seconds, annealing for 20 seconds at 72 °C, extension at 72 °C for 2 minutes 42 seconds, and final extension at 8 minutes for 72 °C was carried out. Then, 1 uL of Dpn1 was added to the reaction and reaction mixture was incubated for 1 hour at 37 °C and stored at 4 °C upon completion. Plasmid fragments were run on a 0.8% agarose gel made using 1X tris acetate-EDTA buffer (TAE). Gel was stained using 50 mL of 1X TAE and 5 uL of 10,000X SYBR Safe stain (Invitrogen). Bands corresponding to the correct fragment were cut out, and fragments extracted using GeneJET gel extraction kit. Purified fragments were combined in an 8:2 molar ratio of fragment 1 (3.1 ng/ul) to fragment 2 (16.6 ng/ul). 10 uL of HiFI DNA assembly master mix (New England BioLabs) was added to the combined fragment solution to get a final volume of 20 uL. This was incubated for 1 hour at 50 °C before being stored on ice and chemically transformed into DH5α cells. The plasmid was then purified from the DH5α cells using a Qiagen QIAprep Spin Miniprep kit and the sequence was confirmed using Sanger sequencing (Genewiz).

### Protein Expression

Untagged wild type GFAP in pET-11a vector codon optimized for E. coli and synthesized by GenScript and C294S GFAP obtained from standard cloning procedures (see previous section) was recombinantly expressed in BL21(DE3) cells. Chemical transformations were performed by incubating BL21(DE3) cells on ice for 15 mins with 1 µl of 0.2 µg/µl plasmid in a standard 1.5 ml microcentrifuge tube. The competent cells were then heat shocked for 90 seconds at 42 °C and then cooled on ice for 2 minutes before the addition of 500 µl of Luria broth (LB) media. This was incubated for 20 minutes at 37 °C with 220 rpm shaking before being plated on an ampicillin LB agar plate. An isolated colony was used to inoculate 500 ml of LB media containing 25 ml of 20% w/v unlabeled glucose, and grown in a gravity incubator overnight at 37 °C. Overnight culture was added to 1 L of LB to an approximate optical density (OD_600_) of 0.01. The cells were grown until an optical density (OD_600_) of 0.8 was obtained (1 cm pathlength, Biochrom Biowave Cell Density Meter CO8). Cells were then harvested by centrifugation at 6,000 g for 10 min before being resuspended in 1 L of M9 minimal media with U-^13^C_6_ glucose and ^15^N ammonium chloride (Cambridge Isotope Laboratories). After 30 min, protein expression was induced by addition of 0.5 M isopropyl-β-D-thiogalactoside (IPTG). The protein was expressed for 3 hr with 220 RPM shaking at 37 °C and the cells harvested by centrifugation at 6,000 g for 10 mins. The cell pellet was flash frozen and stored at −80°C.

### Inclusion Body Purification

The cell pellet was thawed on ice and resuspended in 50 mM tris(hydroxymethyl)aminomethane (Tris-HCl), pH 7.5, 50 mM glucose, 10 mM ethylenediaminetetraacetic acid (EDTA), and 10 mg/mL lysozyme. This mixture was then vortexed for 30 seconds and incubated for 20 min at 37 °C. A solution of 20 mM Tris, pH 7.5, 200 mM sodium chloride (NaCl), 1 mM EDTA, 1% v/v Triton X-100, 10 ug/ ml RNAaseA, 10 µg/ml DNAase1, Magnesium chloride was added to this mixture, which was then incubated for another 20 min at 37 °C and then centrifuged at 5,000 g for 10 min to collect the inclusion body fraction. The supernatant was discarded and the resulting pellet resuspended by pipetting up and down in 10 mM Tris-HCl, pH 7.5, 1 mM EDTA, and 0.5% v/v Triton X-100 and centrifuged at 5,000 g for 10 min. The supernatant was discarded and the resulting pellet resuspended in 10 mM Tris-HCl, pH 7.5, 1 mM EDTA, 1.5 M potassium chloride (KCl) and 0.5% v/v Triton X-100 and centrifuged at 5,000 g for 10 min. The supernatant was discarded and the resultant pellet resuspended in 10 mM Tris, pH 7.5, 1 mM EDTA, 0.15 M KCl, and 0.5% v/v Triton X-100 and centrifuged at 5,000 g for 10 min. The supernatant was discarded and the resultant pellet resuspended in 10 mM Tris-HCl, pH 7.5, 1 mM EDTA, and 0.5% v/v Triton X-100 and centrifuged at 5,000 g for 10 min. Finally, the supernatant was discarded and the resulting pellet resuspended in 10 mM Tris-HCl, pH 7.5, 8 M urea, and 1 mM EDTA and centrifuged at 20,000 g for 10 min. All urea used in the purification and filament preparation steps were deionized using AG® 501-X8 mixed bed resin (Bio-Rad). The supernatant containing the solubilized inclusion bodies was then collected for further purification.

### Affinity Chromatography Purification

The solubilized inclusion bodies were loaded onto a EconoFit Macro-Prep DEAE anion exchange column (Bio-Rad) equilibrated in 10 mM Tris-HCl, pH 7.5, 8 M urea, and 1 mM DTT by hand using a syringe. The protein was carefully eluted using a stepwise gradient ranging from 0 mM NaCl to 500 mM NaCl by hand. Fractions containing GFAP protein were pooled and applied by hand using a syringe onto a EconoFit Macro-Prep High S cation column (Bio-Rad) equilibrated in 10 mM Tris-HCl, pH 7.5, 8 M urea, and 1 mM dithiothreitol (DTT). The protein was then collected in through the flow through, and protein purity confirmed through SDS-PAGE. Purified protein was aliquoted into 1 ml samples, flash frozen and then stored at −80 °C.

### Filament Preparation

Purified GFAP in 10 mM Tris-HCl, pH 7.5, 8 M urea, 1 mM DTT at a concentration of approximately 2.2 mg/ml was dialyzed out of urea in a stepwise fashion to form soluble tetramers. Protein concentration was determined through UV-Vis measurements (Eppendorf BioSpectrometer) using a calculated extinction coefficient of 20,400 M^−1^·cm^−1^ (ProtParam) and theoretical molecular weight of 49.8 kDa. Tetramers were formed using the step-wise removal of urea.^22^ The protein was placed into a 3.5 kDa MWCO dialysis tubing (Spectra Por 3 regenerated cellulose, flat width 29 mm) and dialyzed into 500 ml of buffer containing 10 mM Tris, pH 7.5, 8 M urea, 1 mM DTT, and varying urea concentrations. The dialysis buffer was stirred at room temperature and left in each different urea concentration for 30 min. Urea concentrations were 6 M, 4 M, 2 M, 1 M, and 0 M. The protein was then transferred to 1 L of fresh 0 M urea buffer and dialyzed overnight at 4 °C. Finally, the protein was transferred to fresh 0 M urea buffer and stirred at room temperature for 2 hours to ensure complete removal of urea. To assemble the tetramers into full length filaments, the protein was then dialyzed into an assembly buffer containing 10 mM Tris-HCl, pH 7, 50 mM NaCl, and 1 mM DTT overnight at 29 °C.^23^

### Transmission Electron Microscopy

Filament samples were imaged using Ted Pella ultrathin carbon films on 400 mesh lacey carbon copper grids. TEM grids were prepared following reference 11 by application of 5 µl of protein to the grid surface and incubated for 5 min, then 5 μl of water was then applied and let sit for 1 min before adding 5 μl of 3% uranyl acetate for 3 mins.^23^ Between each step, excess liquid was wicked away using a tissue. The grid was then air dried and imaged on a JEOL-1230 electron microscope with a 2K x 2K Tietz CCD camera operating at 100 keV.

### NMR Measurements

After either fibril or tetramer formation as described above, 8–12 mg of ^13^C and ^15^N labeled protein was collected. The collected protein assemblies were then spun for 3 hours at 233,000 rcf in a swinging bucket rotor. Protein concentration of supernatant was taken to ensure that the majority of protein had been harvested. The resulting filament/tetramer pellet were then transferred to a 3.2 mm thin-walled zirconia ssNMR rotor (Revolution NMR). Before and after protein was added to the rotor, approximately 1 mm PFTE was used as a spacer. To sediment the protein assemblies, the rotor was centrifuged for brief increments at 3,000 rcf. The rotor was spun at 25,000 rcf for two hours to compact the protein sample. Finally, a rotor cap was then attached using cyanoacrylate glue to prevent water loss. Solid state NMR experiments were performed on an 18.8 T Bruker NMR with an Avance III console. A BlackFox NMR and Low-E triple resonance 3.2 mm MAS probe was utilized. Temperature of the sample was kept around 10 ◦ C unless specified otherwise and the spinning rate was 13 kHz.

## Results

### Structural Characterization of Wild-type GFAP Filaments

Fig. 2a shows a negative stain TEM micrograph showing mature GFAP filaments prepared by step-wise removal of chemical denaturant and addition of salt. Magic angle spinning (MAS) ^13^C-^13^C cross polarization-based dipolar assisted rotational resonance (CP-DARR) solid state NMR spectra recorded with high power ^1^H decoupling probe rigid structural motifs in the GFAP filaments. Figure 2b–c shows the carbonyl and aliphatic regions of the CP-DARR spectra recorded from wild-type GFAP filaments. The aliphatic region is shown in Figure 2c, revealing chemical shifts consistent with alpha helical structure, which likely arises from the coiled-coil rod domain of the filament assembly. Signals are observed with NMR chemical shifts consistent with Ile, Ala, Leu, Pro, Glu, Gln, Asp, Thr, and Asn residues. The relatively broad signal intensities observed in Fig. 2b–c may arise from molecular motion or structural heterogeneity within a single filament. However, coiled-coil rod domains of mature intermediate filaments are expected to be rigid and structurally heterogeneous^24^. Given the size of the domain (309 residues) and number of specific amino acids of each type, the relatively broad signal intensities observed for Ala, Ile, Glu, Leu, Asn, Asp, and Val are more likely due to sharp signals from many distinct sites with small differences in their chemical shifts, suggesting a remarkably uniform helical structure. Fig. 2d shows a MAS ^13^C-^13^C INEPT-TOBSY solid state NMR spectrum recorded with weak ^1^H decoupling of GFAP filaments, revealing a small number of highly mobile sites. The observed signals are consistent with random coil chemical shifts for Val, Ile, Leu, Lys, Arg, Gln, Glu, Met, Asp, and Pro residues. These residues likely correspond to the tail domain of GFAP, which has been shown to stick out of the rigid core of filaments formed by the homologous Vimentin protein and is expected to be partially disordered^1,14,25^. To probe regions with motions intermediate to the rigid sites probed by CP-DARR and the highly mobile sites probed by INEPT-TOBSY, we performed ^13^C-^13^C direct polarization DARR (DP-DARR) experiments with high power ^1^H decoupling. Figure 2e shows the difference spectra comparing the ^13^C-^13^C CP-DARR spectrum with the ^13^C-^13^C DP-DARR spectrum. The difference spectrum is mostly devoid of signals, with a few broad and a few sharp signals more prevalent in the CP-DARR spectrum than in the DP-DARR spectrum. These signals are close to the noise level and likely arise from slight differences in experimental conditions. The data are consistent with no significant population of residues with intermediate timescale motions and the spectra in Figure 2 are therefore consistent with significant portions of GFAP in the filaments being rigid and uniformly ordered and a handful of highly mobile disordered residues likely arising from the tail domain.

**Figure 2.**
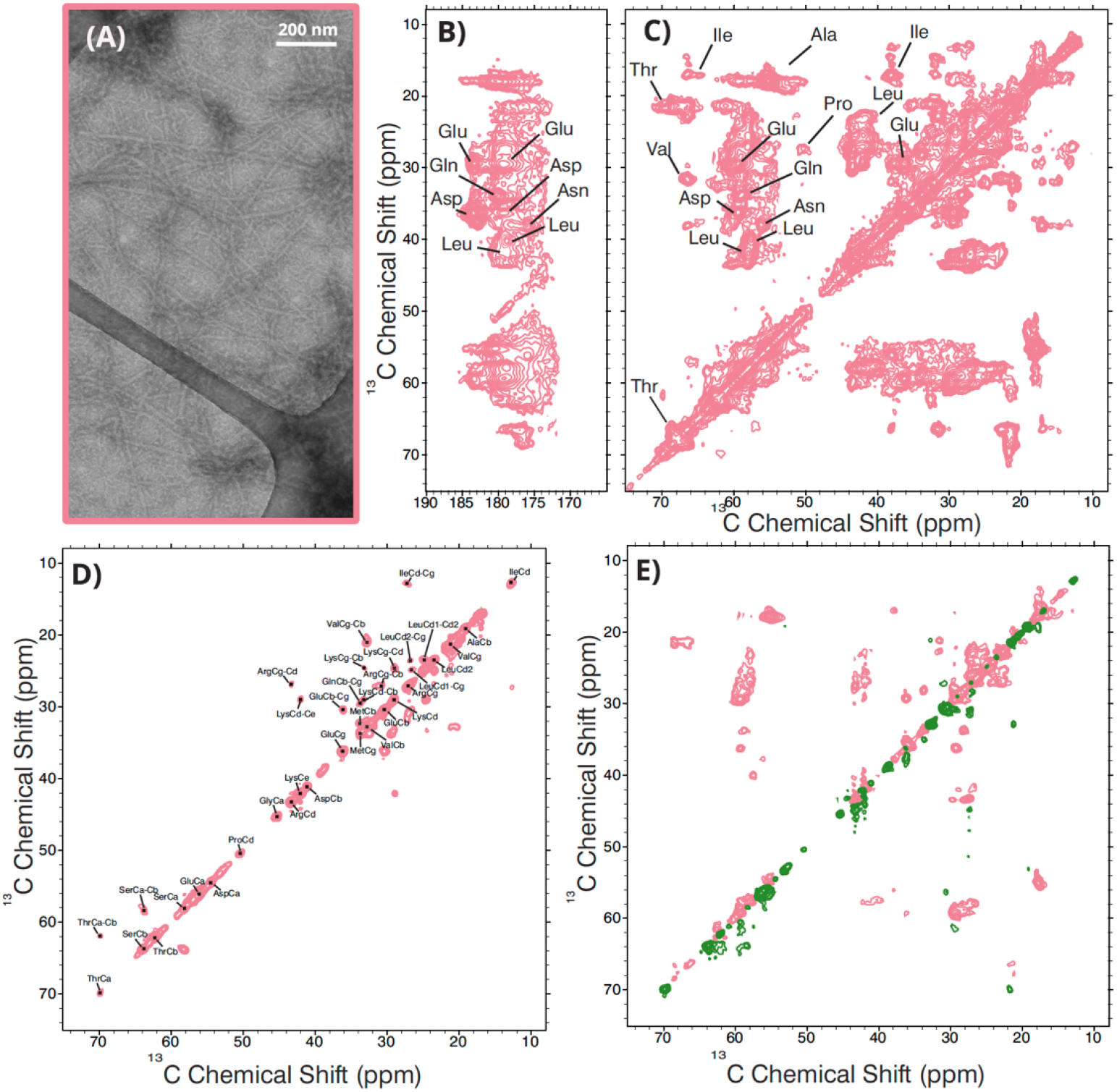
Solid state NMR spectra of wild-type GFAP filaments. (A) Negatively stained TEM image of wild-type GFAP filaments. (B) Carbonyl region of a ^13^C-^13^C CP-DARR spectrum of wild-type GFAP filaments. (C) Aliphatic region of a ^13^C-^13^C CP-DARR spectrum of wild-type GFAP filaments. (D) ^13^C-^13^C INEPT-TOBSY spectrum of wild-type GFAP filaments. (E) Aliphatic regions from a ^13^C-^13^C DARR difference spectrum, obtained by subtracting wild-type filament ^13^C-^13^C CP-DARR spectrum from the wild-type filament ^13^C-^13^C DP-DARR spectrum. Pink contours represent signals that are stronger within the wild-type filament CP-DARR spectrum and green contours represent signals that are stronger in the wild-type filament DP-DARR spectrum.

### Structural Characterization of Wild-Type GFAP Tetramers and Comparison to Wild-Type GFAP Filaments

Removal of denaturant from purified IF proteins under low ionic strength conditions (e.g. no salt) results in tetramers formation. Figure 3a shows negative-stain TEM images for our sample, confirming tetramer formation, which are markedly shorter than the filaments observed in Figure 2a. To probe the rigid structure of the tetrameric assemblies, we recorded ^13^C-^13^C CP DARR spectrum in the same fashion as the filaments. Figure 3b shows the carbonyl region of a difference spectrum obtained by subtracting the wild-type filament ^13^C-^13^C CP DARR spectrum from the wild-type tetramer ^13^C-^13^C CP DARR spectrum. A mixture of sharp and broad or overlapped strong signals corresponding to Ala, Gln, Glu, Lys, Asp, Asn, Ala, Ser, and Thr residues being stronger in the filament spectrum are observed. Figure 3c shows the aliphatic region of the same difference spectrum. Similarly, a mixture of sharp and broad or overlapped strong signals corresponding to Ala, Ile, Leu, Val, Gln, Thr, Glu, Asp, Asn, and Ser residues being stronger in the filament spectrum are observed. Strong signals from the filaments remaining in the difference spectrum are consistent with more residues being ordered in the filaments than tetramers. In support of this interpretation, the ^13^C-^13^C INEPT-TOBSY spectrum of the wild-type tetramers shown in Figure 3d contains many more signals than the same spectrum obtained from the filaments. These additional signals have the random coil chemical shift values expected for Thr, Val, Ala, Lys, Glu, Gln, Met, Asp, and Arg residues, and must arise from highly mobile residues. All peaks observed in INEPT-TOBSY spectrum of the filaments are also observed in the INEPT-TOBSY spectrum of the tetramers (Ile, Val, Leu, Arg, Lys, Glu, Gln, Ser, and Thr residues). Therefore, the residue types observed only in the tetramer spectrum are adopting rigid structure upon conversion to filament structure. Similarly to the filament (Figure 2e), the difference spectrum for the CP-DARR and DP-DARR spectra of the tetramers shown in Figure 3e has weak signals more present in the CP-DARR. The spectrum is therefore confirms that all residues in the tetramers are either highly mobile or very rigid.

**Figure 3.**
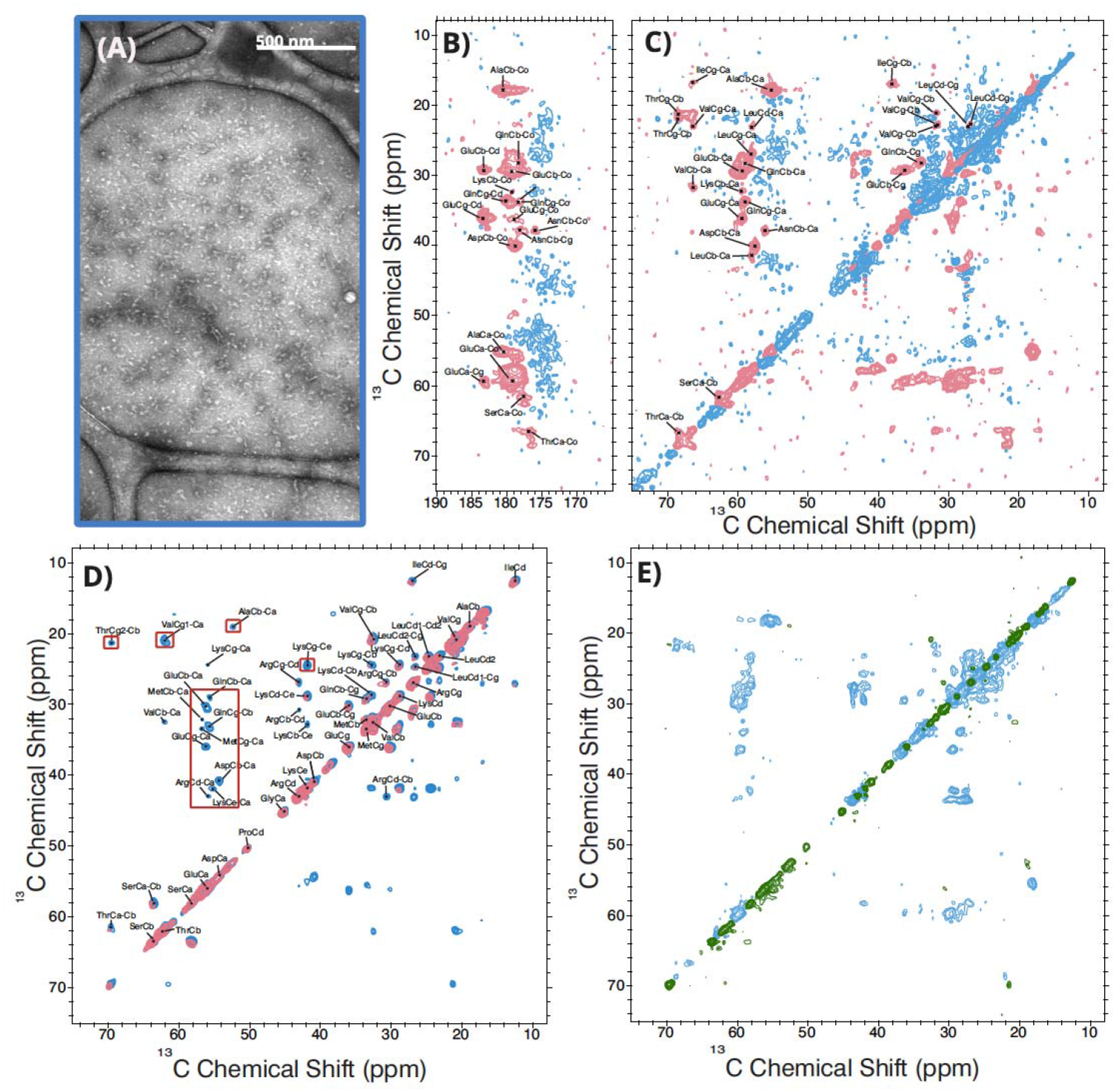
Solid state NMR spectra of wild-type GFAP tetramers. (A) Negatively stained TEM image of wild-type GFAP tetramers. Sections of a ^13^C-^13^C CP-DARR difference spectrum shown in panels (B), carbonyl, and (C), aliphatic, have filament signals in pink and tetramer signals in blue. (D) ^13^C-^13^C INEPT-TOBSY spectrum of GFAP tetramers in blue with the filament spectrum overlayed in pink. (E) Aliphatic region from a ^13^C-^13^C DARR difference spectrum for tetramers comparing CP-DARR in blue and DP-DARR in green.

### Structural Characterization of C294S GFAP Tetramers and Comparison to Wild-Type GFAP Tetramers

The negative-stain TEM image in Figure 4a shows that under the same conditions, the C294S mutant GFAP forms visually similar tetramers to that of wild-type GFAP. Figures 4b–c show carbonyl and aliphatic regions of a difference spectrum calculated from ^13^C-^13^C CP-DARR spectra recorded from C294S mutant and wild-type GFAP tetramers. The spectrum is dominated by noise, consistent with most residues residing in similar conformations and environments in the two samples. A few weak signals are observed in the mutant spectrum corresponding to Glu, Ala, and Gln NMR chemical shifts. The ^13^C-^13^C INEPT-TOBSY spectrum shown in Figure 4d is essentially indistinguishable from the same spectrum recorded from wild-type filaments, showing there is no difference in the highly mobile regions of the tetramer samples. Similarly to Figures 2e and 3e, the difference spectrum calculated from CP-DARR and DP-DARR spectrum recorded on the mutant tetramers shows a few weak signals in the CP-DARR spectrum, consistent with the tetramer assemblies containing only sites of rigid order and significant, disordered, motion.

**Figure 4.**
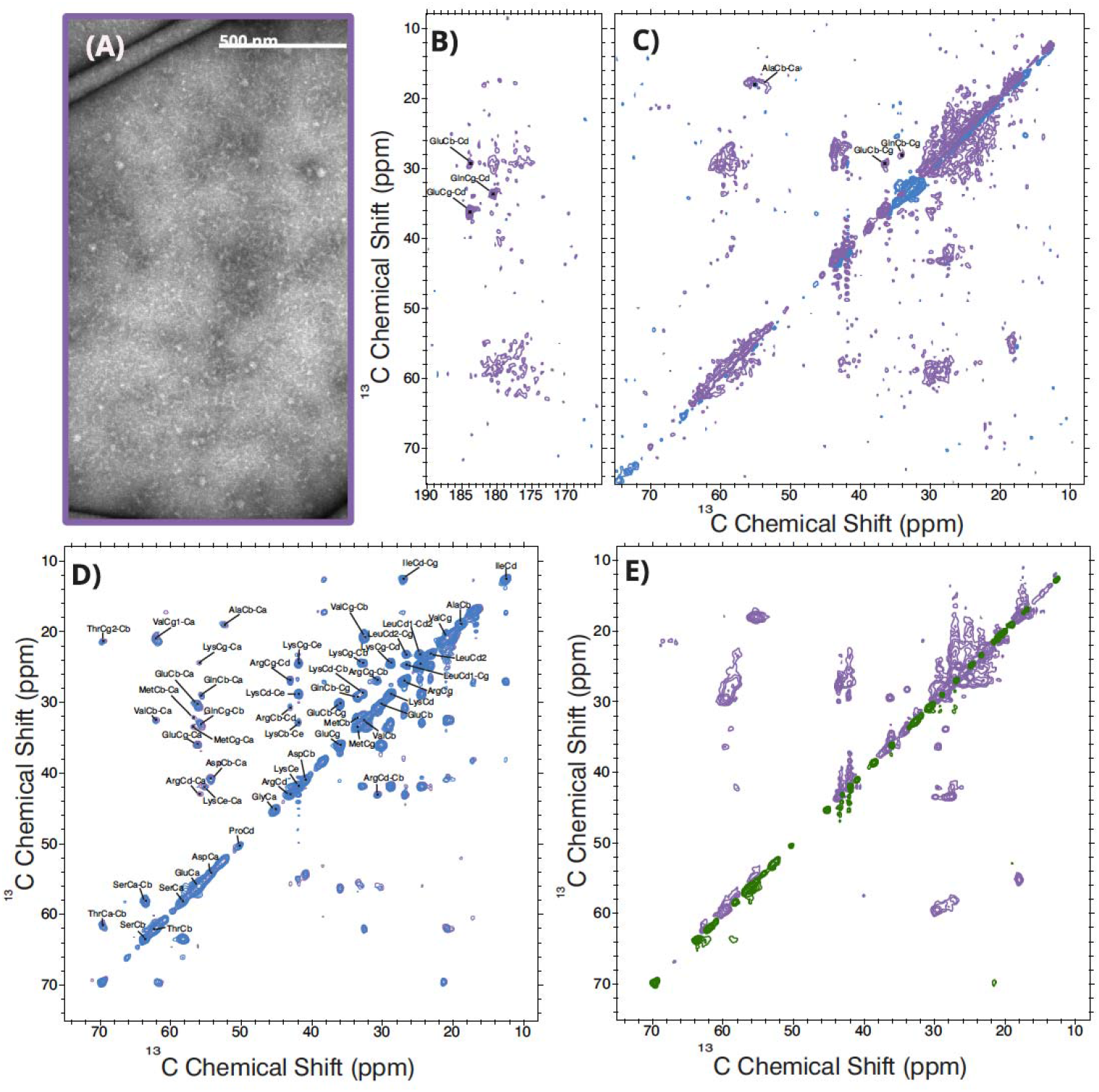
Solid state NMR spectra of C294S GFAP tetramers. (A) Negatively stained TEM image of C294S GFAP tetramers. Sections of a ^13^C-^13^C CP-DARR difference spectrum shown in panels (B), carbonyl, and (C), aliphatic, have wild-type tetramer signals in blue and C294S mutant tetramer signals in purple. (D) ^13^C-^13^C INEPT-TOBSY spectrum of GFAP mutant tetramers in purple with the wild-type tetramer spectrum overlayed in blue. (E) Aliphatic region from a ^13^C-^13^C DARR difference spectrum for mutant tetramers comparing CP-DARR in purple and DP-DARR in green.

### Structural Characterization of C294S mutant GFAP Filaments and Comparison to Wild-Type GFAP Filaments

Unlike C294S tetramers, C294S filaments do not assemble into the expected full-length filaments. The negative-stain TEM images in Figure 5a show C294S mutant GFAP in filament buffer conditions only form short strands that appear to kink and clump together, in stark contrast to the wild-type filaments image in Figure 2a. Additionally, the ^13^C-^13^C CP-DARR difference spectrum calculated from the wild-type filament and mutant filament spectra show significant broad signals in the mutant spectrum not present in the wild-type spectrum. In addition, the wild-type spectrum has a few sharp, strong signals not present in the mutant spectrum. The data are consistent with an increase in heterogenous order in the mutant filament sample compared to the wild-type sample and the loss of a few strongly ordered sites in the mutant sample. Figure 5d shows the ^13^C – ^13^C INEPT-TOBSY spectrum of the mutant filaments has additional signals not present in the same spectrum recorded on the wild-type filaments. Signals corresponding to highly mobile, disordered Glu, Gln, Met, Asp, Lys, and Arg are observed. The INEPT-TOBSY spectrum of the mutant filaments is signficantly more similar to that of the mutant and wild-type tetramers than with the wild-type filaments. The difference spectrum comparting the ^13^C-^13^C CP-DARR and ^13^C-^13^C DP-DARR filament spectra again shows signals more present in the CP-DARR experiment, but is nonetheless consistent with very few residues with motional timescales intermediate to the rigid sites observed in the CP-DARR spectrum and the INEPT-TOBSY spectrum. The signals observed in Figures 5b–d are therefore a characteristic spectroscopic fingerprint for aberrant assembly (i.e. short, kinked filaments) driven by the GFAP C294S mutant GFAP.

**Figure 5.**
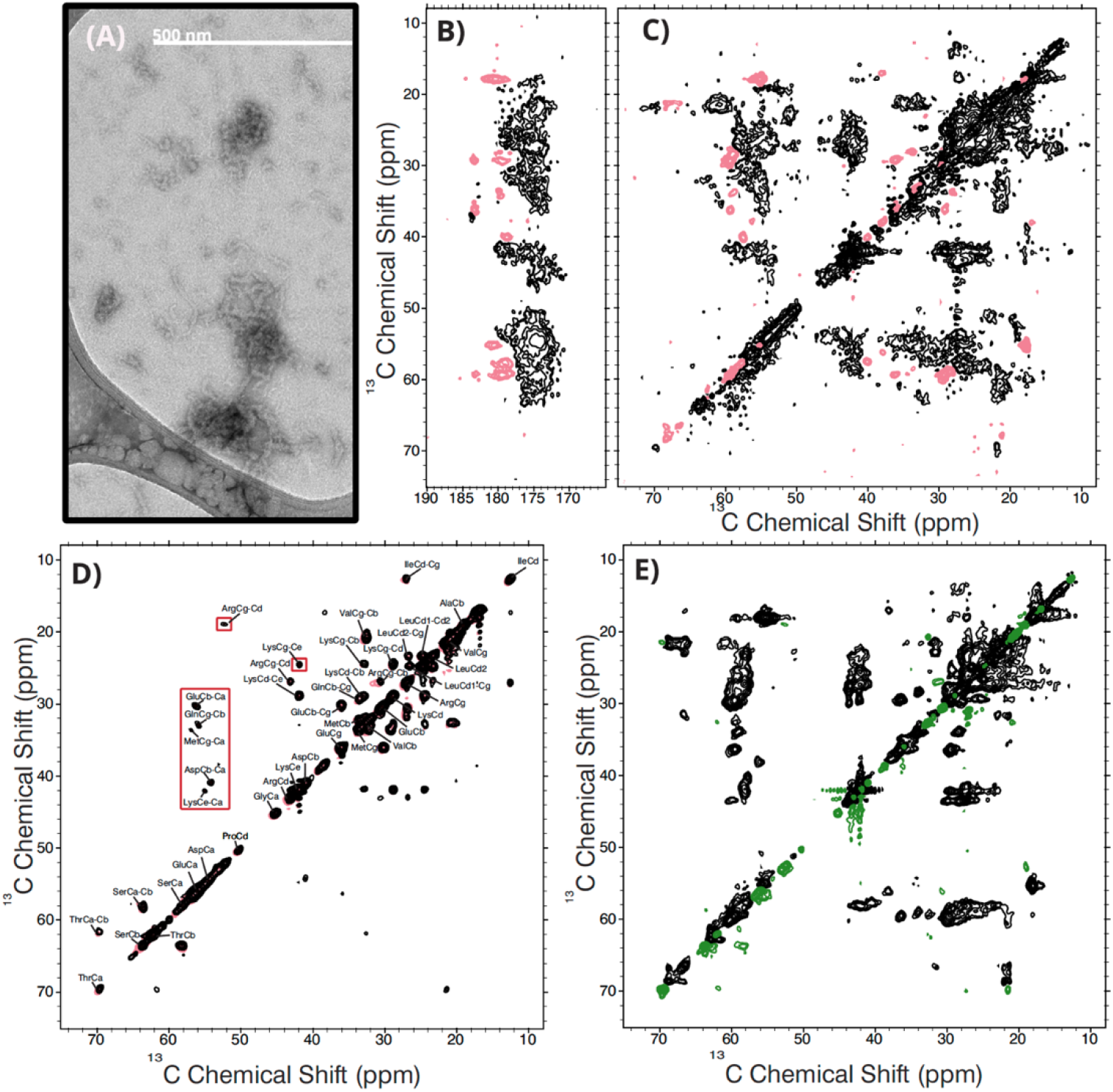
Solid state NMR spectra of C294S GFAP filaments. (A) Negatively stained TEM image of C294S GFAP filaments. Sections of a ^13^C-^13^C CP-DARR difference spectrum shown in panels (B), carbonyl, and (C), aliphatic, have wild-type filament signals in pink and C294S mutant filament signals in brown. (D) ^13^C-^13^C INEPT-TOBSY spectrum of GFAP mutant filaments in brown with the wild-type filament spectrum overlayed in pink. (E) Aliphatic region from a ^13^C-^13^C DARR difference spectrum for mutant filaments comparing CP-DARR in brown and DP-DARR in green.

## Discussion

In summary, our solid state NMR study of GFAP assemblies shows (i) wild-type filaments are primarily rigid with a handful of disordered and highly mobile sites (Figure 2), (ii) wild-type tetramers lose a significant amount of ordered residues coupled with a similar increase in highly mobile and disordered residues (Figure 3), (iii) tetramers of the C294S GFAP mutant are structurally very similar to wild-type tetramers, and (iv) the C294S GFAP mutant does not adopt the conformation of the wild-type filaments and instead more similar, yet not identical, to the wild-type tetramer conformation (Figure 5). Our results are in agreement with previous studies of intermediate filament assemblies showing the intrinsically disordered head and tail domains exposed within the tetramers. Our results are also consistent with the head region becoming more ordered within the filament assembly as a function of end-to-end annealing.

In the homologous vimentin IF protein, absence of the head domain results in inability to assemble.^26^ Furthermore, it has been shown that in vimentin, the last 20 residues of helix 2B found in the rod domain is critical for the tetramer to filament transition.^27^ The highly conserved cysteine is found within those last 20 residues of helix 2B, indicating that structural disruption of helix 2B is what prevents C294S mutant GFAP from assembling into filaments. However, the analogous C328S mutant in vimentin slows down filaments assembly, but does not abolish full-length filaments formation.^28^ Other point mutations at position 328 in vimentin, such as C328A, fail to assemble.^29^ These differential effects of altering the sole cysteine in IF proteins suggest that modification of this site, through oxidation for example, will have differential effects on the various IF proteins and the assemblies they form. In GFAP, as we see that even a conservative mutation of the cysteine to a serine results in the inability of the protein to form filaments, suggesting it may be a more potent sensor of oxidation than vimentin.

## Supplemental Figures

**Figure S1.**
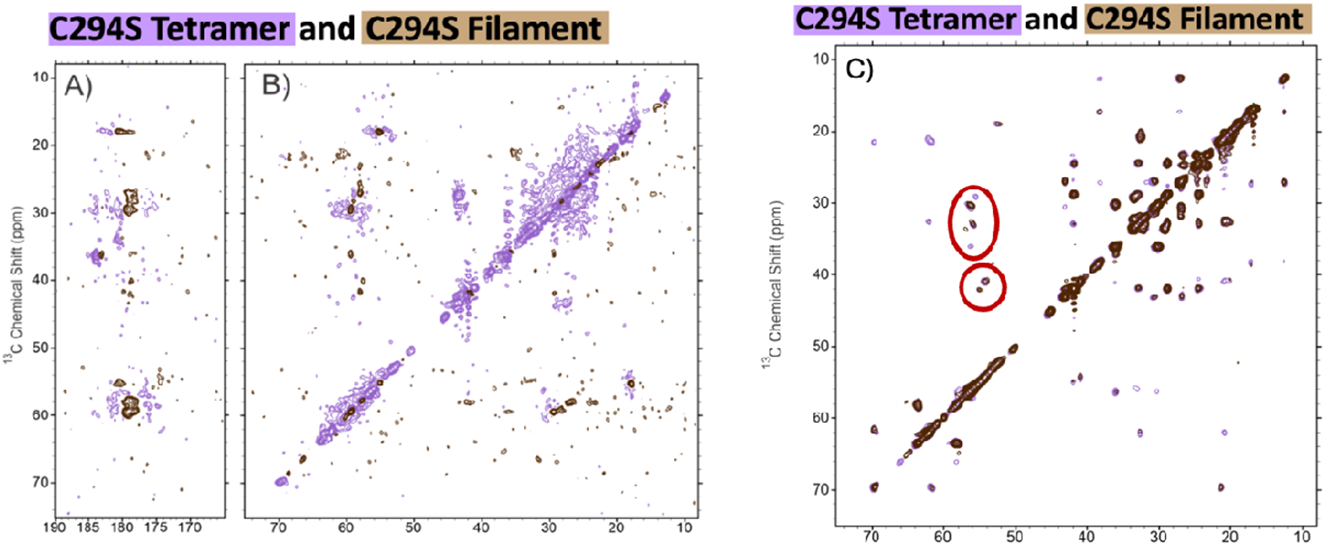
Comparison of C294S Tetramer and Filament Assemblies. A and B show the ^13^C-^13^C CP-DARR difference spectra of the tetramer and filaments. Purple indicates the C294S tetramer spectra, while brown represents the C294S filament signals. A shows the carbonyl region of the ^13^C-^13^C CP-DARR difference spectra and B the aliphatic region. C shows the ^13^C-^13^C INEPT-TOBSY overlay spectra of the C294S tetramer and C294S filament. Peaks absent in the wild-type filament ^13^C-^13^C INEPT-TOBSY spectra that show up in the C294S spectra circled in red.

**Figure S2.**
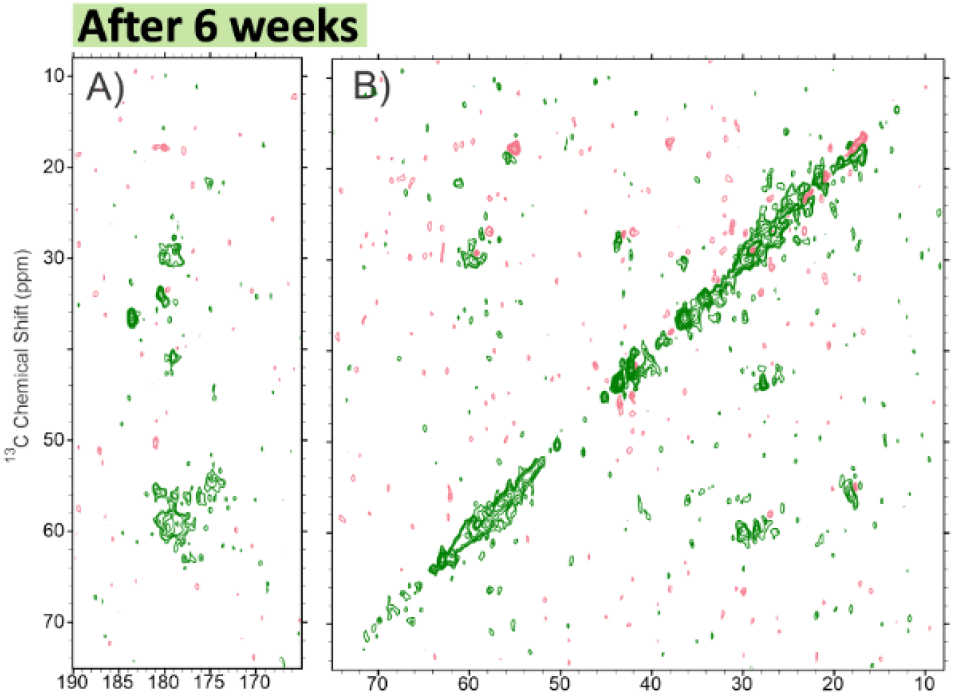
Stability of GFAP Filaments stored in the NMR rotor. A and B show the difference ^13^C-^13^C CP-DARR difference spectra between wild-type filament spectra obtained from the sample NMR sample and stored for six weeks. The signal that corresponds to the freshly taken spectra is in pink and the spectra taken after six weeks is in green. This indicates that the GFAP filaments are stable over time when prepared in the NMR rotor, and that there are no obvious signs of degradation happening within the sample.

## Acknowledgements

The authors would like to thank Dr. John Voss and Dr. Paul Fitzgerald for insightful discussions and helping begin the intermediate filament arm of the Murray Laboratory, and Hillary Sutton for initial work with intermediate filament purification. This work was funded in its entirety by the National Institute of General Medical Sciences of the National Institutes of Health through award R35GM142892 to D.T.M. The content is solely the responsibility of the authors and does not necessarily represent the official views of the National Institutes of Health.

